# Real-time functional connectivity-based neurofeedback of the DLPFC-amygdala pathway during threat-exposure attenuates anxiety

**DOI:** 10.64898/2025.12.22.695740

**Authors:** Wenyi Dong, Stefania Ferraro, Dan Liu, Yuan Zhang, Mengfan Han, Yiqi Mi, Junjie Wang, Kun Fu, Zhiying Zhao, Keith M Kendrick, Dezhong Yao, Shuxia Yao, Benjamin Becker

## Abstract

**Background:** Adaptive regulation of negative emotion is vital for mental health and dysregulations in this domain contribute to the major mental disorders, including anxiety and depression. On the neural level, efficient emotion regulation has been linked to functional communication between the dorsolateral prefrontal cortex (DLPFC) and the amygdala. Gaining voluntary control over this pathway may present an effective strategy for potentiating emotion regulation.

**Objective:** Against this background, we developed a novel connectivity-based real-time fMRI (rt-fMRI) neurofeedback (NF) training that enables individuals to gain volitional control over the DLPFC-amygdala pathway during exposure to threatening stimuli.

**Methods:** The study employed a pre-registered, randomized, sham-controlled design with the experimental group (n = 22) receiving real-time NF information on connectivity between the right DLPFC and the amygdala during threat exposure and the sham control group (n = 23) receiving connectivity NF from a circuit not related to emotion regulation. The ability to maintain regulatory control in the absence of feedback was assessed after four NF runs. Primary outcomes included functional connectivity of the target pathway, as well as anxiety scores.

**Results:** The results demonstrate that successful acquisition of self-regulation of the rDLPFC-amygdala top-down regulatory circuit in the experimental NF group facilitated attenuation of anxiety.

**Conclusion:** In summary, our findings suggest that real-time fMRI neurofeedback (rtfMRI-NF) allows to volitionally enhance rDLPFC-amygdala connectivity, and in turn reduces negative emotional states, rendering the training a promising neurotechnology intervention for mental disorders.

## Introduction

Emotion regulation is important to facilitate adaptive functioning and mental health in our daily life (Côté et al., 2010; Koole, 2009). Deficient emotion regulation represents a risk factor across the major mental disorders (Johnstone and Walter, 2014; Erk et al., 2010; Schulze et al., 2010; Zimmerman et al., 2017; Jiang et al.,2025). Cognitive emotion regulation is one of the most efficient means to regulate negative emotional states and depends on a set of cognitive control functions that enable the selection, implementation, and adjustment of regulation strategies (Egner, 2008). Previous neuroimaging studies point to a key role of the prefrontal cortex (PFC), especially the lateral frontal cortex in cognitive control and emotion regulation (Etkin et al.,2015; Li et al., 2020, Zhou et al., 2019), with the lateral frontal cortex involved in the successful down-regulation of subcortical systems involved in negative emotion reactivity such as the amygdala-centered circuits (Mihov et al., 2013; Gross & John, 2003; Ochsner et al., 2012). In line with this conceptualization studies have reported that the PFC-amygdala pathway plays an important role in emotion regulation (Zhao et al., 2019; Banks et al., 2007; Paret et al., 2016).

Modulation of this pathway has been associated with reductions in anxiety symptoms following behavioural emotion regulation training (Berboth & Morawetz, 2021), real-time functional magnetic resonance imaging neurofeedback (rt-fMRI-NF) training (Zhao et al., 2019), and the anxiolytic effects of pharmacological agents (Dodhia et al., 2014; Gorka et al., 2015) in individuals with heightened anxiety.

Two efficient and extensively studied strategies to attenuate negative emotions are cognitive reappraisal and distraction (Zhao et al., 2021; Kohn et al., 2014; Kanske et al., 2011), which are mainly underpinned by the PFC, particularly the dorsolateral prefrontal cortex (DLPFC) (Cheng et al., 2024; White et al., 2023). In contrast to the ventrolateral prefrontal cortex (VLPFC) consistently engaged in implicit emotion regulation, for instance via its function in selecting goal-appropriate responses and inhibiting inappropriate ones (Braunstein et al., 2017; Zhuang et al., 2021), the DLPFC plays in particular a role in explicit emotional regulation, in particular via orchestrating and regulating other brain regions during this process, including the amygdala, the striatum, the cingulate cortex, and the middle temporal gyrus (MTG) (Northoff et al., 2000; Wager et al., 2008; Zimmermann et al., 2017). Via this function, it may support the reframing of emotional information during reappraisal and re-orienting selective attention during distraction (Liu et al., 2022; Dörfel et al., 2014). The amygdala plays a pivotal role for detecting threats and mediating high-arousing emotions (Frank et al., 2014; Zhang et al., 2025a; Liu et al., 2021), although the effects are to a certain extent depending on functional lateralization (Ocklenburg & Mundorf, 2022). Research has repeatedly shown altered amygdala engagement during cognitive emotion regulation in patients with mental disorders, including patients with depression or addiction (Erk et al., 2010; Zimmerman et al., 2017; Zilverstand et al., 2017). In healthy subjects, increases in functional connectivity (FC) between the amygdala and DLPFC have been reported during successful regulation of emotional responses to threatening stimuli using reappraisal strategies (e.g., detachment) (Erk et al., 2010; Morawetz et al., 2017) or distraction strategies (Kanske et al., 2011), with the strength of DLPFC-amygdala FC being positively correlated with emotion regulation success (Morawetz et al., 2017).

Given the essential role of the DLPFC and the bilateral amygdala, and particularly their dynamic interplay, in emotion regulation, developing modulation approaches targeting the DLPFC-amygdala circuit may not only enhance our understanding of the mechanisms underlying emotion regulation but also bring new possibilities for improving clinical symptoms associated with emotion regulation impairments. Data from the World Health Organization (WHO) delineates the tremendous personal and socio-economic costs of mental disorders and that approximately one-third of the patients do not adequately respond to first-line treatments, including pharmacotherapy like antidepressants or behavioral interventions (Misaki et al., 2024). This underscores the necessity of developing new approaches to complement traditional interventions. Real-time functional magnetic resonance imaging neurofeedback is an innovative neuromodulation technique that allows individuals to gain voluntary control over regional brain activity or connectivity (Kvamme et al., 2022; Linhartova et al., 2019; Zhang et al., 2025b). With its noninvasive nature and whole-brain coverage, a growing body of translational research highlighted the clinical potential of rt-fMRI NF in symptom alleviation in patients suffering from depression, addiction, anxiety, and trauma-related disorders (Gu et al.,2025; Taebi et al.,2024; Misaki et al., 2024; Pindi et al., 2022; Dudek & Dodell-Feder, 2021; Zweerings et al,2020). These findings suggest that rt-fMRI NF may serve as an effective complement to traditional psychotherapeutic approaches.

The majority of previous studies have concentrated on region-based rt-fMRI-NF, particularly involving the amygdala, prefrontal cortex (PFC), insula (INS), and have shown that the regulatory control can be maintained in the absence of feedback (Yao et al., 2016; Zhang et al., 2023). More specifically, rt-fMRI-NF research demonstrated that neurofeedback training effectively downregulates amygdala activity in response to negative stimuli, compared to training without feedback (Brühl et al., 2014; Herwig et al., 2019; Watve et al., 2024).

Furthermore, it was shown that participants were able to enhance anterior insula (AI) activity, accompanied by increased empathic responses to painful stimuli (Yao et al., 2016). Rt-fMRI-NF induced greater AI activity positively correlated with increased interoceptive accuracy after training (Zhang et al., 2023), thereby aiding participants in regulating emotions through interoceptive strategies. In posttraumatic stress disorder, successful downregulation of bilateral amygdala activity via rt-fMRI NF was accompanied by increased activation in both the DLPFC and VLPFC and increased connectivity of the amygdala-PFC pathway (Nicholson et al., 2017).

Collectively, these findings indicate that rt-fMRI NF, based on these regional signals, is a promising therapeutic potential in facilitating emotion regulation. However, there is still a lack of evidence whether direct modulation of the DLPFC-amygdala pathway based on rt-fMRI NF training can also improving emotion regulation.

Initial studies have employed rt-fMRI NF training based on FC during which participants learn to regulate the functional interaction (connectivity) between two brain regions. For instance, participants receiving rt-fMRI-NF focused on the DLPFC-anterior cingulate cortex (ACC) circuit demonstrated increased FC within this circuit, which was associated with reduced anxiety levels (Morgenroth et al., 2020). Additionally, a study by Zhao et al. (2019) found that participants who engaged in active neurofeedback training were able to enhance FC between the right VLPFC (rVLPFC) and the amygdala, leading to decreased anxiety levels in participants with high trait anxiety. These studies highlighted the potential of FC-based neurofeedback training to guide enhanced emotion regulation.

Although the DLPFC is crucial for emotion regulation, the degree to which FC between the DLPFC and the amygdala can be modulated through rt-fMRI-NF training, as well as the varying effects of such modulation on negative emotions, remains to be explored. The present study therefore aims to bridge this gap by exploring whether rt-fMRI NF training targeting the DLPFC and bilateral amygdala pathway can modulate FC between these regions and in turn improve the regulation of negative emotions.

## Materials and methods

### 2.1. Participants

For this randomized double-blind study, 50 healthy participants (25 males, mean age = 22.2 years, SD = 1.74) were recruited from the University of Electronic Science and Technology of China (UESTC). This sample size was determined using the G*Power v3.7 toolbox to detect effects in a mixed ANOVA design, with a statistical power greater than 0.80 (effect size = 0.25, α = 0.05). The estimated sample size was consistent with previous rt-fMRI-NF studies (Linhartova et al., 2019; Zhang et al., 2023) and adequate to identify reliable effects of neurofeedback training at both behavioral and neural levels. Prior to the enrollment of the experiment, a short interview was conducted to exclude any previous or current psychiatric, neurological or major health disorders. A total of 25 participants (13 males) were assigned to the experimental (EXP) group who received NF based on the FC between the bilatAMY and the right DLPFC (rDLPFC). In contrast, 25 participants (13 males) were assigned to the sham control (SHAM) group, receiving sham NF derived from the FC between the bilateral primary motor cortex (bilatM1) and right Heschl’s gyrus (rHES). This sham condition has been shown to effectively control for major confounding factors such as motivation, placebo effects, and non-specific global activations (Zhao et al., 2019; Wang et al., 2010). Five participants were excluded from the analysis due to incomplete participation (n = 2) or excessive head movement (n = 3). Consequently, the final sample included 22 participants (10 males; age, M ± SD = 22.2 ± 2.02 years) in the EXP group, and 23 participants (11 males; age, M ± SD = 22.3 ± 1.48 years) in the SHAM group.

All procedures of the current study were approved by the local ethical committee at UESTC and followed the latest version of the Declaration of Helsinki. All participants provided informed consent. This study was preregistered on clinicaltrials.gov (https://ichgcp.net/zh/clinical-trials-registry/NCT06033053). Participants completed all tasks during a single experimental session. Prior to MRI acquisition, participants received a comprehensive overview of the procedure, which included pre- and post-MRI assessments, a practice session for familiarizing with the fMRI task, and the rt-fMRI-NF session comprising the functional localizer task (LocT), 4 runs of NF training task (NFT), and the transfer task (TransT). Participants were instructed to minimize head movement during scanning and were informed that they could withdraw from the experiment at any time and for any reason.

### 2.2. Experimental protocol

#### 2.2.1 Behavioral assessment and statistics

To ensure the comparability of the two groups in terms of personality and emotional functioning, all participants completed the State-Trait Anxiety Inventory (STAI) (Spielberger et al., 1971), the Beck Depression Inventory (BDI) (Beck et al., 1996), the Emotion Regulation Questionnaire (ERQ) (Gross & John, 2003) before the MRI scan. After each NFT run, participants reported the strategies they had found effective (see Table S2 for details). Following the entire MRI scanning, the STAI-S was retested.

Group differences for demographic variables and questionnaire scores were tested using independent t-tests. While differences between pre- and post-NFT tests of STAI-S scores were computed with a repeated-measures ANOVA, in which the TIMEPOINT (pre, post) was used as the within-subject factor and the GROUP (EXP and SHAM) was used as the between-subject factor. A repeated measures ANOVA was carried out on the self-reported effectiveness of the emotion regulation strategy (hereafter, strategy validity ratings) obtained after each of the NFT run (see the next section), in which the RUN (run1, run 2, run3, run 4) was used as the within-subject factor and GROUP (EXP and SHAM) was used as the between-subject factor.

#### 2.2.2 rt-fMRI-NF instructions

Participants were informed that the rt-fMRI-NF task was designed to improve their emotion regulation skills, enabling them to improve their ability to cope with negative emotional events in daily life and reducing stress. The two groups (i.e., EXP and SHAM) were identically instructed to explore and identify effective strategies for emotion regulation during NFT when threatening picture stimuli were displayed, with the aim of increasing the thermometer bars. As no specific strategy is required for successful NF training (Thibault et al., 2018; Sepulveda et al., 2016; Zhao et al., 2019), participants were not provided with predefined strategies to regulate their negative emotions. Thus, they were encouraged to find their own effective approaches to manage the negative emotions and to regulate the level of the feedback bars (see Fig.1B) without using of physical means such as changing their breathing or body movements to influence the bar level. Participants were told that although the feedback (i.e., the level of the thermometer bar) was computed in real-time, it would represent brain connectivity with an approximate delay of about 8 seconds for technical reasons. They were also told that, at the end of each NFT run, they would be asked (through a rating scale) whether they had identified an effective strategy for raising the level of the thermometer bar (see Table S1) and that, once such a strategy had been identified, they would have to apply it consistently in the remaining NFT and TransT runs. After the experimental overview, a shorter version of the LocT was practiced outside the scanner to ensure proper understanding of the task before proceeding with the MRI acquisition.

**Fig. 1.**
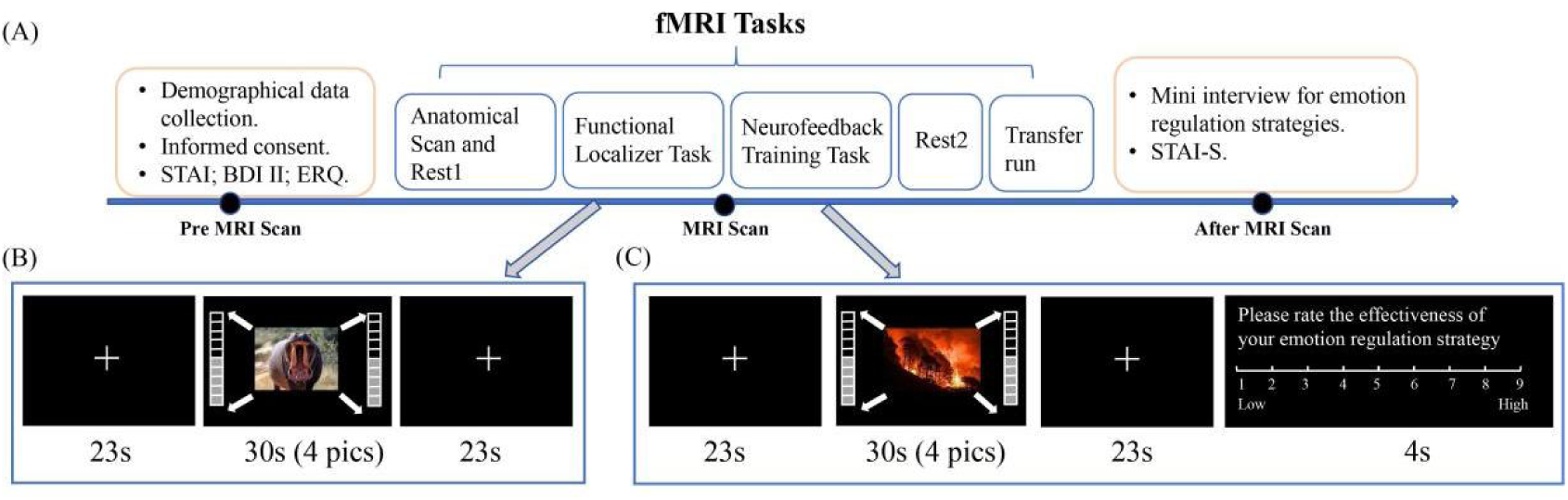
Experimental Procedures (A) Timeline of the experimental procedures for both the EXP and SHAM groups. The whole procedure lasted about 120 minutes. (B) Representation of the functional localizer task. (C) Representation of the neurofeedback training task. EXP: experiment, SHAM: sham control

#### 2.2.3 Definition of regions of interest (ROIs)

The training circuit for the EXP group was defined individually for each subject and was based on regions of interest (ROIs) localized in the rDLPFC (functionally defined) and bilatAMY. Based on the contrast between threatening stimuli and fixation cross during the LocT, we identified, in an online manner, the peak activity within a mask comprising the right frontal pole and right middle frontal gyrus, as defined by the Harvard-Oxford atlas (see Fig. S1A). At this peak, we created a 6-mm spherical ROI for the rDLPFC (see Table S1). The bilatAMY ROI was created by combining 2 masks located in the right (rAMY, MNI coordinate at [22_-3_-18]) and left amygdala (lAMY, MNI coordinate at: [-23_-4_-18]). The rDLPFC mask and the bilatAMY ROI were derived from the Harvard-Oxford atlas. The training circuit for the SHAM group was constituted by a ROI defined in the rHES (MNI coordinate at [48_-14_4] obtained from the anatomy toolbox’s TE1.0 region) and by a ROI defined by 2 masks located in the bilateral M1 (bilatM1, MNI coordinate at [± 38_-22_56]) (Wang et al., 2010) (See Fig. S1B for two training circuits). For both groups, a white matter ROI (MNI coordinate at [21_-32_-36]) (Zhao et al. 2019) was used to regress out the white matter signal during the real-time calculation of the FC (i.e., NFT). All the ROIs were created as spheres of 6-mm radius.

#### 2.2.4 rt-fMRI-NF experiment

##### Localizer task (LocT)

During the LocT, participants were asked to view carefully the presented images and not use any strategies to control their emotions. This task consisted of 6 blocks and each block lasted 30 seconds. During each block, 4 threatening picture stimuli (valence: M ± SD = 3.03 ± 0.77; arousal: M ± SD = 6.04 ± 0.65) were presented.

Consistent with Zhao et al. (2019), each stimulus was displayed for 7.5s, gradually increasing in size from half to full size, to incrementally increase the perceived threat. A 23s fixation cross interval was used between regulation blocks.

##### Neurofeedback training task (NFT)

The 4 NFT runs followed the same protocol to the LocT run except that threatening pictures (valence: M ± SD = 3.06 ± 0.84; arousal: M ± SD = 6.03 ± 0.61) were presented alongside with a real-time NF visualized as a thermometer on either side of the screen (see Fig.1) reflecting the real-time FC of the selected training circuit (rDLPFC-bilatAMY for the EXP group or rHES-bilatM1 for the SHAM group). During the presentation of the threatening pictures, participants were asked to downregulate the level of the thermometer using the discovered strategy (please see the rt-fMRI-NF instructions paragraph).

After each NFT run, participants were asked to rate their perceived effectiveness of the strategy on a 1-9 scale (strategy validity ratings; 1 = not effective at all; 9 = very effective) in changing the bar level of the thermometer within 4s (see Fig.1). The thermometer representing the FC values was updated every 2s.

##### Transfer task (TransT)

Following the 4 NFT runs, a TransT run was conducted to assess whether participants could maintain their ability to regulate the FC of the selected circuit without NF. This TransT run mirrored the NFT runs, with the only difference being that participants were provided with no real-time NF information but instead a thermometer with fixed bars. Participants were instructed to continuously use the regulation strategy they found most effective during the NFT and also to provide their ratings on the strategy effectiveness at the end of the transfer run.

### 2.3. Image data acquisition

MRI images were collected using a 3T, GE Discovery MR750 scanner (General Electric Medical System, Milwaukee, WI, USA). High resolution whole-brain volume T1-weighted images were firstly collected with a 3D spoiled gradient echo pulse sequence (TR: 6 ms; TE: minimum; flip angle, 12°; FOV: 256 × 256 mm; voxel size = 1×1×1 mm; 172 slices). Functional images in all the tasks (i.e., the LocT, NFT, and TransT) were acquired using a T2*-weighted EPI sequence (TR = 2000 ms; TE = 30 ms; flip angle = 90°; FOV = 220 × 220 mm²; matrix = 64 × 64; 32 slices; slice thickness = 3.4 mm; gap = 0.6 mm). Pre- and post-training resting-state fMRI data were also collected but not reported in this study.

### 2.4. fMRI data analyses

#### 2.4.1. Real-time NF analyses

The Turbo-BrainVoyager version 4.2 toolbox (Brain Innovation, Maastricht, The Netherlands) was used during the MRI acquisition to process both the T1-weighted structural image and the real-time functional data (LocT and NFT runs). The rt-fMRI-NF setup employed in the present study was in line with procedures used in our earlier research (Zhao et al., 2019). Specifically, brain images were transmitted in real-time from the MRI scanner to the local disk of the TBV-installed computer. After preprocessing and normalization of the T1-weighted images, the functional data were overlaid onto the processed and coregistered anatomical image in native space to ensure accurate localization of the functionally defined ROI (i.e., rDLPFC). ROIs initially defined in MNI space were transformed back into native space using the inverse normalization parameters obtained during the T1 normalization process. Real-time FC for the selected target circuit (rDLPFC-bilatAMY for the EXP condition or rHES-bilatM1 for the SHAM condition) was computed as the partial correlation between time series of the selected ROIs with a default window length of 20 time points by including time series from the white matter ROI as a covariate, while correcting for head movements.

#### 2.4.2. Offline task-evoked activation analyses

The SPM12 software (Wellcome Department of Cognitive Neurology, London, UK; http://www.fil.ion.ucl.ac.uk/spm) was employed for offline fMRI data processing to identify brain regions involved in the processing of threatening images or emotion regulation in the LocT, NFT, and TransT. After excluding the first 5 volumes, imaging data of each run underwent canonical preprocessing comprising head-motion correction, co-registration of the mean functional image with the T1-weighted image, normalization to the standard MNI space, and smoothing (8 mm full-width at half-maximum Gaussian kernel). For both groups (EXP and SHAM), 3 first-level generalized linear models (GLMs) were constructed for the LocT, NFT, and TransT runs, respectively. While the GLM design matrix of the LocT run included one regressor (threatening stimuli blocks), the NFT runs, and the TransT included two regressors (regulation during threatening stimuli blocks and regulation rating at the end of each run), all convolved with the canonical hemodynamic response function. All design matrices also included the six head motion parameters as nuisance regressors.

To identify brain regions involved in different tasks, contrast images in the LocT from each subject in the EXP or SHAM group were subjected to the group-level analysis in each group separately using a one-sample t-test in SPM12. Similarly, contrast images of the regulation block from each group were analyzed separately using one-sample t-tests in the NFT or TransT. Group differences were examined by using independent t-tests. At the whole-brain level, results were corrected using a *p* < 0.01 FDR-corrected threshold and only clusters larger than 10 voxels were reported (see Fig S2 for details).

#### 2.4.3. Offline functional connectivity analyses

To examine changes in FC between the target ROIs during NF trianing, a generalized psychophysiological interaction analysis (gPPI) was conducted using the CONN FC toolbox v22 (www.nitrc.org/projects/conn). The default pipeline preprocessing and denoising steps were firstly performed (Nieto, 2020). These preprocessing steps generated several covariates that were included in the first level analysis as nuisance variables (‘scrubbing’, containing the scans identified as outliers for each subject/session; ‘QC_timeseries’ containing the timeseries of the global signal changes and framewise displacement; ‘realignment’ containing six rigid-body parameters representing subject motion). Then, in the experimental set-up, the regulation condition was explicitly modeled by entering block onset times and duration.

Whole-brain seed-based gPPI analyses were then performed for different tasks (i.e., the LocT, NFT and TranT) by computing bivariate correlations between the rDLPFC seed and all brain voxels during regulation (Nieto et al., 2020). The seed region was defined as a 6-mm sphere centered on the peak activation (MNI coordinates: 54, 26, 29) within the rDLPFC mask in the LocT of both the EXP and SHAM groups. Second-level analyses were then performed in SPM12 to characterize the seed-based whole-brain FC patterns within each group (*p* < 0.01, voxel-level FDR-corrected) and between-group comparisons for NFT runs (*p* < 0.01, FDR-corrected).

To specifically test FC differences between the EXP and SHAM groups in the rDLPFC-bilatAMY circuit in different tasks, we imported produced maps (i.e., whole-brain rDLPFC seed-based map) into MarsBar (Brett et al., 2002) to extract beta values from the bilatAMY ROI of each subject representing the FC strength of the rDLPFC-bilatAMY circuit. The beta values from the LocT and TranT runs were entered into two separate independent t-tests with one assessing potential group differences prior to NFT (LocT) and the other evaluating the maintenance effect of NF training (TranT). The FC values from the NFT runs, were instead analyzed using a repeated-measures ANOVA with RUN (run1, run 2, run3, run 4) as the within-subject factor and GROUP (EXP and SHAM) as the between-subject factor. In addition, the FC values from the TransT run were compared with the mean of four NFT runs to further assess the maintenance of training success in both the EXP and SHAM groups.

## Results

### 3.1. Demographics and behavioural results

Two-sample t-tests on demographics (age) and questionnaire scores (i.e., BDI, STAI-T, and ERQ) revealed no significant differences between the EXP group and SHAM group (*all ps* ≥ 0.1; see Table 1). For pre–post comparisons of state anxiety changes between the two groups, a repeated-measures ANOVA on STAI-S scores showed a marginal main effect of GROUP (F _1,43_ = 3.918, *p =* 0.054, Ƞ ^2^ = 0.084), with a higher level of state anxiety in the SHAM compared with the EXP group. However, no significant main effect of TIMEPOINT (F _1,43_ = 1.282, *p* = 0.264, Ƞ*_p_* ^2^ = 0.029) or GROUP × TIMEPOINT interaction (F _1,43_ = 1.370, *p* = 0.248, Ƞ*_p_* ^2^ = 0.031) were found. To exclude the possibility of significant group difference at pre-training, pairwise comparisons were conducted: pre-training STAI-S (*p* = 0.207; Table1 and Fig. 2A) did not show any significance difference between the two groups, while there was a marginally higher level of state anxiety in the SHAM than the EXP group at post-training (*p* = 0.052). These results suggest that the marginal main effect of group was mainly driven by an increase of anxiety following the NFT task in the SHAM group, which was not observed in the EXP group.

**Fig. 2.**
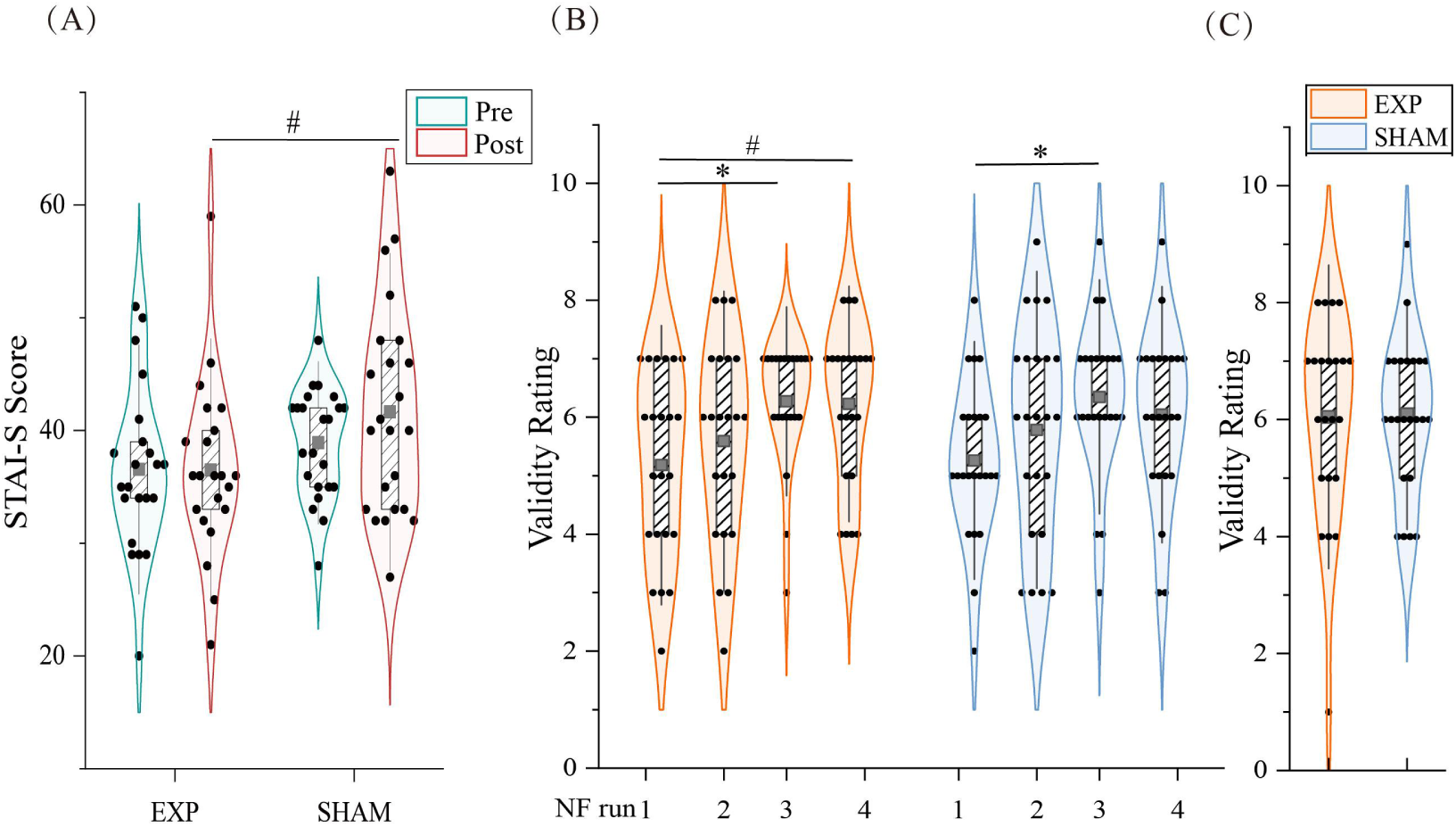
Behavioral results (A) STAI-S scores for the EXP and SHAM groups in pre- and post-test (# *p* = 0.052). (B) Validity ratings of emotion regulation strategies following each NFT run in the EXP and SHAM groups. (C) Validity ratings of emotion regulation strategies following the TransT run in EXP and SHAM groups. Error bars represent standard error and gray rectangular represents the mean value (*: *p* < 0.05, # *p* < 0.1)

**Table 1.** Demographic and questionnaires statistics in the EXP and SHAM groups before and after the MRI scan (M ± SD).

| Measurements | EXP | SHAM | t-value | p-value |
| --- | --- | --- | --- | --- |
| Age (year) | 22.2 ± 2.02 | 22.30 ± 1.48 | 0.133 | 0.895 |
| STAI-State_pre | 36.6 ± 7.37 | 38.9 ± 4.81 | 1.281 | 0.207 |
| STAI-State_post | 36.5 ± 7.77 | 41.65 ± 7.39 | 2.001 | 0.052 |
| STAI-Trait | 39.1 ± 6.82 | 39.4 ± 6.79 | 0.192 | 0.849 |
| BDI | 26.3 ± 5.01 | 26.9 ± 5.98 | 0.334 | 0.740 |
| ERQ | 45.2 ± 6.17 | 47.6 ± 6.52 | 1.258 | 0.215 |

\_pre: pre-test; \_post, post-test. STAI: State-Trait Anxiety Inventory;
BDI: Beck Depression inventory; ERQ: Emotion Regulation Questionnaire.

A repeated-measures ANOVA was conducted on effectiveness rating scores of the emotion regulation strategy (i.e., strategy validity ratings) during NFT runs and revealed a significant main effect of RUN (F _3,129_ = 7.772, *p <* 0.001, Ƞ ^2^ = 0.153). Post-hoc analyses showed a significant increase from run 1 to run 3 (*p* < 0.001) and run 4 (*p* = 0.005; Fig. 2B). However, there were no significant main effect of GROUP (F _1,43_ = 0.016, *p =* 0.900, Ƞ*_p_* ^2^ = 0.000) or GROUP × RUN interaction (F _3,129_ = 0.206, *p =* 0.892, Ƞ*_p_* ^2^ = 0.005). These results suggest a general increase in the feelings of effectiveness of using regulation strategies over NFT runs in both the EXP and SHAM groups. In the TransT, the group difference was not significant (t _1,43_ = 0.091, *p* = 0.928, Cohen’s d = 0.027; Fig. 2C).

### 3.2 Neurofeedback training (NFT) effects on changes in functional connectivity

Changes in FC of the target circuit over runs in different tasks were presented in Fig. 3A. In the LocT, the two groups did not show significant differences in the extracted FC strengths of the rDLPFC-bilatAMY pathway (t _1,43_ = 0.581, *p* = 0.565, Cohen’s d = 0.175; two-tailed), suggesting no significant difference between the two groups before NFT.

**Fig. 3.**
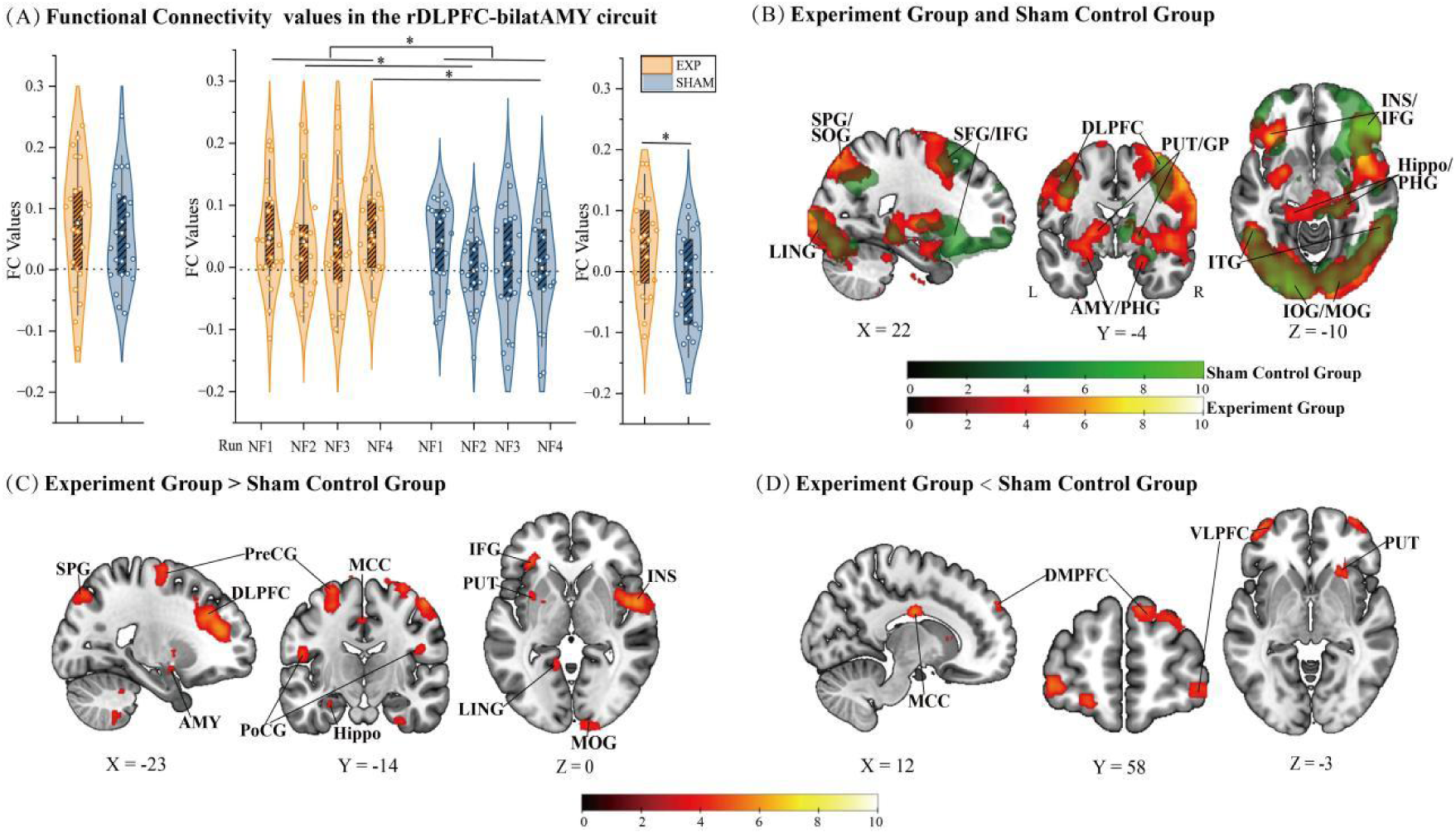
Functional connectivity results (A) Violin plot of FC values in the rDLPFC-bilatAMY circuit for the EXP and SHAM groups in the LocT, NFT and TranT runs respectively. (B) Whole-brain functional connectivity based on the rDLPFC (a 6-mm sphere ROI centered on [54, 26, 29]) during NFT runs in the EXP group (red) and the SHAM group (green). (C) Between-group comparison (EXP > SHAM) for the whole brain level functional connectivity (FC) based on the rDLPFC during NFT runs (*p* < 0.01, FDR-corrected). (D) Between-group comparison (EXP < SHAM) for the whole brain level FC based on the rDLPFC during NFT runs (*p* < 0.01, FDR-corrected). LING: lingual gyrus; SPG: superior parietal gyrus; SOG: superior occipital gyrus; ITG: inferior temporal gyrus; MCC: middle cingulate gyrus;. Hippo: hippocampus gyrus; PHG: parahippocampal gyrus; AMY: amygdala; INS: insula; IFG: inferior frontal gyrus; PUT: putamen; GP: globus pallidus; IOG: inferior occipital gyrus; MOG: middle occipital gyrus; DMPFC: dorsomedial prefrontal cortex; VLPFC: ventrolateral prefrontal cortex,( *: *p* < 0.05). Error bars represent standard error, white rectangular represents the mean value

During the 4 NFT runs, the repeated-measures ANOVA on the extracted FC values from the rDLPFC-bilatAMY circuit showed a significant main effect of GROUP (F _1,43_ = 5.578, *p* = 0.023, Ƞ ^2^ = 0.115) indicating of stronger rDLPFC-bilatAMY FC in the EXP compared with the SHAM group. Neither the main effect of RUN (F _3,129_ = 0.936, *p* = 0.424, Ƞ ^2^ = 0.021) nor the GROUP × RUN interaction effect reached significance (F _3,129_ = 0.841, *p* = 0.474, Ƞ ^2^ = 0.019). Exploratory post-hoc tests showed that the EXP group exhibited significantly stronger FC of the rDLPFC-bilatAMY circuit compared to the SHAM group in run 2 (*p* = 0.036) and run 4 (*p* = 0.019) NFT runs.

In the TransT, an independent t-test on FC values of the rDLPFC-bilatAMY circuit revealed a significant group difference between the two groups (t_1,43_ = 2,689, *p* = 0.01, Cohen’s d = 0.802; two-tailed), as reflected by greater connectivity in the target pathway in the EXP group as compared to the SHAM group in the absence of feedback. To further examine the maintenance of training efficacy, we also compared the mean of rDLPFC-bilatAMY FC values across the 4 NFT runs with FC values in the TransT run in the EXP group. A paired-t test revealed no significant differences in this comparison within the EXP group (t _21_ = 0.248, *p* = 0.807, Cohen’s d = 0.132; two-tailed). These findings suggest that successful regulation of the target pathway via NF training can be maintained even without feedback.

To provide a more complete view of NF-induced FC changes beyond the target pathway, we reported the whole-brain level FC pattern of the seed rDLPFC and found similar patterns between the two groups in the NFT. More specifically, both groups showed enhanced FC of the rDLPFC with brain regions in (i) the cognitive control and emotional regulation networks including the bilateral DLPFC, bilateral inferior prefrontal gyrus (IFG) and bilateral insula (INS), (ii) the learning and memory regions including the bilateral hippocampus (Hippo) and parahippocampal gyrus (PHG), (iii) the visual information processing and integration regions including the bilateral lingual gyrus (LING), bilateral inferior occipital gyrus (IOG) extending to the bilateral middle occipital gyrus (MOG) the bilateral superior occipital gyrus (SOG), and the bilateral inferior temporal gyrus (ITG) (*p* < 0.01, FDR-corrected; Fig. 3B). Furthermore, the EXP group demonstrated FC additionally with the bilatAMY and PUT extending to the GP (*p* < 0.01, FDR-corrected). Importantly, further between-group comparisons of the rDLPFC-seeded whole-brain FC showed that the EXP group exhibited stronger connectivity with the left DLPFC (lDLPFC), PUT, amygdala, hippocampus, PHG, middle cingulate gyrus (MCC), LING and the right STG, INS and MOG (EXP > SHAM; Fig. 3C and Table 2), but decreased connectivity with the bilateral VLPFC and the right dorsomedial prefrontal cortex (DMPFC), dorsal ACC (dACC), and IFG (EXP < SHAM; Fig. 3D and Table 2).

**Table 2.** Between-group comparisons of the whole-brain level functional connectivity based on the rDLPFC.

| Brain region | BA | No. Voxels | Peak t-value | X | Y | Z |
| --- | --- | --- | --- | --- | --- | --- |
| **EXP > SHAM** |  |  |  |  |  |  |
| L. Dorsolateral Prefrontal Cortex | 47/9/46 | 1830 | 6.69 | -28 | 30 | 28 |
| Dorsolateral Prefrontal Cortex |  |  | 6.42 | -26 | 40 | 16 |
| Inferior Frontal Gyrus |  |  | 5.42 | -38 | 26 | 8 |
| R. Superior Temporal Gyrus | 22/6/21 | 1233 | 6.37 | 48 | 0 | -10 |
| Superior Temporal Gyrus |  |  | 5.99 | 54 | -2 | 0 |
| Insula |  |  | 5.83 | 46 | 2 | 0 |
| R. Precentral Gyrus | 6/3/4 | 731 | 6.22 | 58 | -6 | 48 |
| Precentral Gyrus |  |  | 5.9 | 52 | -14 | 60 |
| Postcentral Gyrus |  |  | 4.26 | 48 | -14 | 24 |
| L. Superior Parietal Gyrus | 7 | 255 | 5.69 | -18 | -80 | 48 |
| Superior Parietal Gyrus |  |  | 4.89 | -24 | -74 | 46 |
| R. Inferior Frontal Operculum | 9/45 | 90 | 5.65 | 60 | 16 | 28 |
| Superior Temporal Gyrus |  |  | 3.96 | 66 | 0 | 8 |
| Inferior Frontal Gyrus |  |  | 3.74 | 58 | 26 | 22 |
| L. Superior Frontal Gyrus | 6 | 402 | 5.63 | -26 | -10 | 68 |
| Precentral Gyrus |  |  | 5.10 | -26 | -10 | 56 |
| Precentral Gyrus |  |  | 4.47 | -26 | -20 | 70 |
| R. Superior Frontal Gyrus | 6 | 207 | 5.53 | 32 | -10 | 70 |
| Precentral Gyrus |  |  | 4.6 | 24 | -16 | 78 |
| R. Cuneus | 7/19 | 166 | 5.46 | 12 | -86 | 46 |
| Superior Parietal Gyrus |  |  | 5.39 | 20 | -84 | 50 |
| Superior Parietal Gyrus |  |  | 4.45 | 18 | -74 | 62 |
| R. Calcarine | 18 | 158 | 5.11 | 14 | -102 | -4 |
| Middle Occipital Gyrus |  |  | 4.25 | 22 | -102 | -2 |
| L. ParaHippocampal | 36/20 | 156 | 4.84 | -30 | -20 | -22 |
| Hippocampal |  |  | 4.39 | -28 | -18 | -20 |
| Fusiform |  |  | 3.86 | -30 | -36 | -20 |
| L. Precentral Gyrus | 43 | 171 | 4.82 | -46 | -2 | 22 |
| Postcentral Gyrus |  |  | 4.68 | -52 | -14 | 18 |
| Rolandic Operculum |  |  | 4.08 | -46 | -8 | 14 |
| L. Lingual Gyrus | 19/29 | 99 | 4.63 | -12 | -50 | -4 |
| Posterior Cingulate |  |  | 3.77 | -12 | -52 | 6 |
| R. Fusiform | 20 | 60 | 4.51 | 30 | -10 | -34 |
| ParaHippocampal |  |  | 3.92 | 28 | -2 | -32 |
| L. Putamen |  | 127 | 4.33 | -26 | -2 | -10 |
| Amygdala |  |  | 4.29 | -24 | -2 | -12 |
| Putamen |  |  | 3.91 | -22 | 0 | 2 |
| Putamen |  |  | 3.82 | -32 | 6 | 0 |
| L. Middle Cingulate Cortex | 31 | 24 | 4.02 | -2 | -14 | 46 |
| R. Superior Temporal Pole | 38 | 30 | 4.29 | 38 | 14 | -26 |
| R. Cerebellum Anterior Lobe |  | 33 | 4.39 | 8 | -48 | -22 |
| L. Cerebellum Anterior Lobe |  | 31 | 4.08 | -24 | -42 | -32 |
| R. Cerebellum Posterior Lobe |  | 50 | 4.57 | 26 | -46 | -48 |
| L. Cerebellum Posterior Lobe |  | 545 | 5.08 | -18 | -48 | -50 |
| Cerebellum Posterior Lobe |  |  | 4.94 | -26 | -52 | -52 |

**b)**
| Brain region | BA | No. Voxels | Peak t-value | X | Y | Z |
| --- | --- | --- | --- | --- | --- | --- |
| **SHAM > EXP** |  |  |  |  |  |  |
| R. Angular | 7/40 | 361 | 6.82 | 34 | -58 | 38 |
| R. Superior Frontal Gyrus | 6/8/9 | 138 | 5.75 | 8 | 34 | 44 |
| Superior Frontal Gyrus |  |  | 4.28 | 4 | 38 | 54 |
| Dorsal Anterior Cingulate Cortex |  |  | 4.23 | 6 | 36 | 30 |
| L. Ventrolateral Prefrontal Cortex | 10 | 272 | 5.66 | -46 | 52 | -6 |
| Ventrolateral Prefrontal Cortex |  |  | 5.57 | -40 | 58 | -4 |
| Superior Frontal Gyrus |  |  | 5.11 | -18 | 60 | -14 |
| R. Inferior Frontal Operculum | 9 | 298 | 5.62 | 36 | 12 | 28 |
| Dorsomedial Prefrontal Cortex |  |  | 5.02 | 36 | 16 | 38 |
| R. Lateral Ventricle | 40 | 113 | 5.53 | 4 | 18 | 4 |
| S. Dorsomedial Prefrontal Cortex | 10 | 118 | 5.46 | 8 | 58 | 36 |
| Superior Frontal Gyrus |  |  | 4.59 | 28 | 56 | 30 |
| Superior Frontal Gyrus |  |  | 4.39 | 20 | 60 | 30 |
| R. Superior Frontal Gyrus | 10 | 185 | 5.38 | 36 | 64 | 0 |
| Ventrolateral Prefrontal Cortex |  |  | 4.89 | 46 | 56 | -8 |
| Ventrolateral Prefrontal Cortex |  |  | 4.61 | 38 | 60 | -10 |
| R. Superior Frontal Gyrus | 8 | 221 | 5.36 | 38 | 22 | 58 |
| Superior Frontal Gyrus |  |  | 5.08 | 28 | 26 | 60 |
| Superior Frontal Gyrus |  |  | 4.46 | 24 | 32 | 54 |
| R. Inferior Frontal Gyrus | 47 | 256 | 5.23 | 28 | 34 | -8 |
| Putamen |  |  | 5.07 | 22 | 22 | -2 |
| Medial Orbitofrontal Cortex |  |  | 4.74 | 22 | 40 | -14 |
| R. Supramarginal Gyrus | 40/2 | 328 | 5.08 | 50 | -42 | 44 |
| Inferior Parietal Gyrus |  |  | 4.57 | 44 | -52 | 52 |
| Supramarginal Gyrus |  |  | 4.17 | 54 | -32 | 38 |
| L. Inferior Temporal Gyrus |  | 45 | 5.01 | -44 | -26 | -26 |
| R. Inferior Temporal Gyrus |  | 89 | 4.78 | 54 | -34 | -18 |
| Inferior Temporal Gyrus |  |  | 4.09 | 48 | -28 | -22 |

All results were reported with a threshold of $p < 0.01$ FDR-corrected.

For report completeness, we also examined task-evoked activation at the whole brain level in the three tasks, see supplementary materials (Fig. S2) for details.

## Discussion

The present study investigated the effectiveness of a connectivity-based rt-fMRI-NF approach for enhancing emotion regulation and reducing anxiety. Feedback from the rDLPFC-bilatAMY pathway, yet not the sham motor control pathway-allowed subjects in the EXP group to acquire rapid control over the connectivity of the target pathway across four training runs, with significantly stronger connectivity in the EXP compared with the SHAM group. These findings align with previous research (e.g., Zhao et al., 2019; Yao et al., 2016; Robineau et al., 2017) reporting that individuals can learn volitional control over brain function by means of regional activity- or connectivity-informed NF. Furthermore, the EXP group maintained their regulation ability in a TransT run without NF assistance. Behaviorally, NF training prevented an increase of anxiety levels evoked by threatening stimuli during NF training runs, as reflected by a trend of anxiety increase following the NFT in the SHAM relative to the EXP group.

During emotional processing before the NFT - that is in the activation during the LocT run - the two groups exhibited comparable FC in the rDLPFC-bilatAMY circuit. However, when subjects were provided with real-time NF, they exhibited a better ability in volitional control over the FC of the target circuit, as reflected by a significantly stronger FC in the EXP compared with the SHAM group, particularly in the second and the fourth training runs. FC changes of the rDLPFC-bilatAMY pathway were similar to our previous study targeting on the VLPFC-amygdala pathway (Zhao et al., 2019). The strongest FC values of the circuit observed in the fourth run may suggest a cumulative learning process with the progress of the four NFT runs (Watanabe et al., 2017; Zhao et al., 2019). Consistent with previous studies based on either regional activity or FC-based NF (Yao et al., 2016; Zhang et al., 2023, 2025; Zhao et al., 2019), we also found that subjects in the EXP group could maintain their ability in increasing FC of the target pathway, as supported by a significantly stronger FC in the EXP than the SHAM group and an insignificant difference between the mean FC across the 4 NF runs and the TransT run. These findings on the one hand demonstrate again the feasibility of achieving volitional control over the top-down emotion regulation pathway and are also important to translational application of rt-fMRI NF on the other, particularly in terms of the maintenance effect indicative of the possibility of eliminating sustained dependence on the expensive MRI scanner.

At the behavioral level, we found a trend of increases in anxiety levels following the NFT in the SHAM compared with the EXP group, which could be induced by exposure to threatening stimuli during NFT in the SHAM group and be eliminated by successful up-regulation of the rDLPFC-bilatAMY connectivity in the EXP group. However, although the EXP group exhibited superior performance in regulating the target pathway, the two groups showed comparable feelings of effectiveness of using regulation strategies (i.e., validity ratings) in the NFT. Given that successful regulation was accompanied by higher validity ratings when a specific strategy was provided (Zhang et al., 2023), we speculated that the use of free strategies in the present study – requiring trial and error to identify the optimal strategy – may have blunted group differences in subjective ratings of strategy effectiveness.

This progressive increased confidence of applying strategies over training runs did not necessarily translate to enhanced neuromodulation, which highlights the critical role of real-time NF in acquiring self-regulation ability and validates the blinding procedure to participants.

For FC changes beyond the target pathway, the EXP and SHAM groups showed similar FC patterns between the rDLPFC and brain regions in the emotional regulation (e.g., DLPFC, IFG and INS; Banks et al., 2007; Berboth et al., 2021; Dörfel et al., 2014), learning and memory (e.g., hippocampus and parahippocampal gyrus; Horga et al., 2015; Murty et al., 2010), and visual information processing and integration systems (e.g., IOG, SOG, LING and ITG; Gholam & Daliri., 2025; Yang et al., 2015; Kang et al., 2011). Given the difference of whether receiving real NF between the two groups, these similar FC patterns may therefore suggest shared neural mechanisms underpinning processing of threatening stimuli, regulation strategy application and evaluation of synchronization between strategy application and NF. Importantly, compared with the SHAM group, the EXP group showed stronger FC between the rDLPFC and the left DLPFC and PUT, amygdala, hippocampus, LING, IFG, and the right INS and MOG, a set of regions associated with emotional regulation and processing (Banks et al., 2007; Berboth et al., 2021; Dörfel et al., 2014), learning and memory (Horga et al., 2015; Murty et al., 2010), and visual processing (Gholam & Daliri., 2025; Yang et al., 2015; Kang et al., 2011). While enhanced connectivity with the lDLPFC and IFG may reflect congenerous emotion regulation processing with the seed rDLPFC, increased connectivity with emotional processing regions (i.e., amygdala and INS) further confirmed findings based on analysis of extracted FC values of the target pathway and may indicate a better regulatory effect on negative emotional responses in the EXP group. For increased FC with regions in the learning/memory and visual processing systems, they may reflect better learned associations between regulation strategy application and NF, consistent with the perspective that NF training is a special form of reinforcement learning (Lubianiker et al., 2022), and enhanced visual input of NF information matching regulation effort in the EXP group. By contrast, the SHAM group exhibited stronger FC with the VLPFC, DMPFC, dACC, and IFG, a set of regions mainly engaged in cognitive control and conflict monitoring (Vermeylen et al., 2020; Oehrn et al., 2014; Kim et al., 2013) and therefore suggest that subjects in the SHAM group may experience conflict between regulation effort and incongruent NF, which also requires a higher level of cognitive control during such less effective regulation.

In addition to the NF-induced changes in FC, we also examined task-evoked brain responses. Similar to the comparable FC patterns of the target pathway, comparable patterns of task-evoked brain responses were also found in the LocT, including brain regions from distributive networks engaged in emotional regulation (e.g., DLPFC, IFG and bilatAMY; Etkin et al., 2015; Frank et al., 2014; Kanske et al., 2011), learning and memory (e.g., hippocampus, PUT, and GP; Patterson & Knowlton., 2018; Tricomi et al., 2009; Seger., 2006), and visual information processing and integration (e.g., IOG, MOG and MTG;Yang et al., 2015; Nasaruddin et al., 2014; Park et al., 2010). These results suggest similar neural processes between the two groups when applying strategies for emotion regulation in response to threatening stimuli prior to receiving NF training. In the NFT, while the two groups showed generally similar brain response patterns, except in the TransT, where significant activity in the amygdala was only found in the SHAM group. This is in line with the finding of more effective regulation of the rDLPFC-bilatAMY connectivity in the EXP group during NFT and previous findings of decreased amygdala activity but increased FC of amygdala with the DLPFC during emotion regulation when exposed to threatening stimuli (Erk et al., 2010; Kanske et al., 2011; Morawetz et al., 2017). The absence of significant amygdala (and the INS) reactivity in the EXP group during the TransT, suggesting a maintenance effect of NF training on inhibiting negative emotional reactivity to threatening stimuli. Of note, although these effects align with significant group differences of NF-induced FC changes of the target circuit, we did not observe significant group differences for these task-evoked brain responses. They are therefore more at the descriptive level and should be regarded as supplementary support for the significant NF effect on the target circuit.

Findings of the present study need to be discussed in the context of limitations. Firstly, the present study recruited healthy subjects and we only used the STAI-S to measure NF-induced changes in anxiety. Although the STAI is one of the most commonly used questionnaires for anxiety, it may not be sensitive enough to measure NF-induced changes in anxiety particularly in healthy population. Secondly, we only found a marginal NF effect on anxiety, although we obtained a significant effect on the neural level via one visit of NF training. While there is ample evidence for rapid acquirement of volitional control over brain activity or connectivity after brief NF training (e.g., Yao et al., 2016; Zhang et al., 2023, 2025b; Zhao et al., 2019), more robust and long-lasting effects on behavioral or symptomatic improvement are of greater significance for translational application. Future studies are needed to optimize NF training protocol such as dividing multiple training sessions in different visits for most robust training effects at both the behavioral and neural levels. Finally, given that no specific strategy is required for successful NF training (Thibault et al., 2018; Sepulveda et al., 2016; Zhao et al., 2019) and to keep consistent with our previous study (Zhao et al., 2019), we did not provide instructed strategies to subjects for regulating the target pathway. Although the relative superiority of specific vs. free strategies remains controversial, it is possible that neural modulation of the rDLPFC-bilatAMY pathway, which is engaged in explicit top-down emotion regulation, may be more effective using specific strategies (e.g., reappraisal or distraction). Future studies are also needed in this context.

In conclusion, the present randomized, double-blind, sham-controlled study provides evidence for the efficacy of rt-fMRI NF training in modulating FC of the DLPFC-amygdala pathway involved in explicit emotion regulation. Successful regulation of this pathway can be maintained in a TransT run without NF information and induces a trend of decreased anxiety in participants receiving real-time NF of the DLPFC-amygdala connectivity compared with those receiving sham NF. Beyond the target pathway, NF-induced changes in FC are also found in emotional regulation, learning and memory, and visual information processing and integration systems reflective of coordination across multiple brain systems for supporting successful neural modulation based on FC-based NF training. Taken together, these findings suggest the translational potential of FC-based NF training targeting on emotion regulation circuits in improving anxiety, although further optimization is still needed, particularly in terms of achieving more robust behavioral and symptomatic changes.

## Supporting information

Supplemental Tables and Figures will be used for the link to the file on the preprint site

## Acknowledgments

This work was supported by the STI 2030-Major Projects (grant number: 2022ZD0208500), the National Natural Science Foundation of China (grant number: 82271583) and the National Natural Science Foundation of China (grant number: 32471139).

## Author contributions

B.B. S.Y. S.F., and W.D. designed the study; W.D., S.F., D.L., Y.Z., M.H. and Y.M. conducted the experiment and collected the data; W.D., Y.Z., J.W. and K.F. performed the data analysis; W.D. and S.F. wrote the manuscript draft; K.K., D.Y., S.Y. and B.B. critically revised the manuscript draft.

