## Supplemental Tables and Figures will be used for the link to the file on the preprint site for "Real-time functional connectivity-based neurofeedback of the DLPFC-amygdala pathway during threat-exposure attenuates anxiety"

**Table S1.**

Center MNI coordinates of the experiment participants’ right DLPFC ROIs during NFT. All ROIs were defined during the localizer run (LocT) and constructed as spheres with a radius of 6 mm. .

| **EXP**  **group** | **X** | **Y** | **Z** |
| --- | --- | --- | --- |
| s1 | 46 | 27 | 28 |
| s2 | 38 | -1 | 47 |
| s3 | 42 | 3 | 41 |
| s4 | 38 | -1 | 46 |
| s5 | 43 | 0 | 43 |
| s6 | 49 | 4 | 42 |
| s7 | 29 | 46 | 25 |
| s8 | 45 | 3 | 34 |
| s9 | 46 | 40 | 8 |
| s10 | 29 | 21 | 44 |
| s11 | 35 | 24 | 43 |
| s12 | 37 | 30 | 42 |
| s13 | 42 | 15 | 48 |
| s14 | 48 | 11 | 42 |
| s15 | 36 | 25 | 41 |
| s16 | 38 | 13 | 45 |
| s17 | 41 | 18 | 46 |
| s18 | 50 | 23 | 40 |
| s19 | 40 | 27 | 37 |
| s20 | 31 | 30 | 42 |
| s21 | 37 | 32 | 43 |
| s22 | 40 | 23 | 38 |

**Table S2.**

Emotion regulation strategies used in NFT runs and TransT run.

**a)**

| **EXP**  **group** | **Gender** | **Strategies** |
| --- | --- | --- |
| s1 | Male | Visualize traveling with my girlfriend, feeling completely relaxed. |
| s2 | Male | Reducing focus on gory elements diminishes their psychological impact. |
| s3 | Female | Imagine having abundant wealth to spend, creating joyful mental states. |
| s4 | Female | Silently recite classical poems/song lyrics internally. |
| s5 | Female | Recall humorous moments and happy memories with friends. |
| s6 | Female | Avoid viewing bloody visuals, obscuring them reduces perceived threat. |
| s7 | Female | Think about loved ones and cherish shared positive experiences. |
| s8 | Male | Replay beautiful memories or imagine vibrant fantasy scenarios. |
| s9 | Female | Reinterpret scenes through sci-fi lenses or idealized childlike perspectives. |
| s10 | Female | Immersing in the scene and  thinking solutions. |
| s11 | Male | Analyzing scenarios objectively with intellectual pride. |
| s12 | Male | Redirect attention away from negative thought patterns. |
| s13 | Male | Enter a meditative state of mental emptiness. |
| s14 | Male | Focus on counting while viewing  the image. |
| s15 | Male | Reimagine spiders/snakes as affectionate personal pets. |
| s16 | Male | Disregard disturbing imagery while contemplating unrelated relaxing topics. |
| s17 | Female | Stare at neutral areas while cognitively reframing them as non-threatening. |
| s18 | Female | Analyze details calmly while letting the mind wander freely. |
| s19 | Female | Pretend to converse with others or self-reflect – this can promote relaxation. |
| s20 | Female | Mentally mock the image through humorous internal commentary. |
| s21 | Female | Visualize cathartic emotional release (e.g., screaming). |
| s22 | Male | Imaging the stimuli not so threatening, and reappraisal. |

**b)**

| **SHAM group** | **Gender** | **Strategies** |
| --- | --- | --- |
| s1 | Male | Focus on non-central parts of the image and associate  them with  other objects or scenarios. |
| s2 | Male | Imagine it is a movie scene: immerse myself while adding imagined details. |
| s3 | Male | Stare at less frightening areas and  transform them into pleasant imagery. |
| s4 | Male | Combine my personal experiences with the experimental materials to explore  creative possibilities. |
| s5 | Female | Convince myself the horror elements couldn't happen in reality while focusing onnon-threatening areas. |
| s6 | Male | Develop solutions to overcome fear, or consciously avoid what can't be resolved. |
| s7 | Female | Maintain relaxation, consciously associating with opposite/positive scenarios. |
| s8 | Female | Gradually regulate emotions by avoiding the most intense visual stimuli. |
| s9 | Female | Suppress negative emotions, emphasizing positive feelings. |
| s10 | Female | Divert attention completely away from threatening parts. |
| s11 | Female | Visualize positive memories or mentally replay calming music. |
| s12 | Male | I immerse myself in the scene as if I were physically present. |
| s13 | Female | Observe details and imagine elements appear in real world. |
| s14 | Male | Mentally devise prevention strategies and reframe perspectives. |
| s15 | Male | Roleplay as either participant or observer through self-questioning. |
| s16 | Female | Hyper-focus on the scene while imagining actively experiencing the process. |
| s17 | Male | Mentally position myself as the victim in the scenario. |
| s18 | Male | Engage in self-immersion to consciously track emotional responses. |
| s19 | Female | Intentionally focus on the most colorful elements of the image. |
| s20 | Female | Observe while associating the scene with unrelated non-threatening contexts. |
| s21 | Female | Adopt first-person perspective to think responsive  actions. |
| s22 | Female | Reframe violent/graphic contents as fictional representations. |
| s23 | Male | Restrict visual attention to minimally disturbing areas. |

###### Selection of the negative stimuli

Prior to the rt-fMRI experiment, 48 participants (16 females, 32 males; mean age = 21.3 years, SD = 1.92) were recruited to rate 144 emotional images. Of these, 39 images were selected from the International Affective Picture System (IAPS; Lang et al., 2008) and were also used in a previous study by Zhao et al. (2019), while the remaining 115 images were downloaded from the internet. All images were broadly categorized into six types: body, animal, blood, fire, people, and object.

During the rating task, participants were first asked to select an emotion word (fear, disgust, or joy) that best described their response to each image. They then rated the valence of each image on a 9-point scale (1 = extremely negative, 9 = extremely positive) and the arousal level on the same scale, with higher scores indicating stronger emotional responses. Mean valence and arousal ratings were subsequently calculated for each image. Images with mean valence ratings of approximately 3 and mean arousal ratings of approximately 6 were selected as stimuli for each run of the fMRI tasks, ensuring a balanced distribution of valence, arousal, and negative stimulus categories.

**Table S3.**

Arousal rating, valence rating, the rate and the stimuli distribution in six categories of the selected negative stimuli during locT run, four NFT runs and the Transfer run.

|  | **Arousal** | | **valence** | | **Rate** | | **Stimuli Distribution in Four Categories** |
| --- | --- | --- | --- | --- | --- | --- | --- |
|  | Mean | SD | Mean | SD | Mean | SD |  |
| **Localizer** | 6.04 | 0.65 | 3.03 | 0.77 | 0.69 | 0.09 | Body: 6; Animal: 8; Blood: 5; Object: 1; Fire: 4 |
| **NFT run1** | 6.04 | 0.66 | 2.93 | 0.84 | 0.65 | 0.10 | Body: 8; Animal: 8; Blood: 3; Object: 1; Fire: 4 |
| **NFT run2** | 6.04 | 0.70 | 3.16 | 0.90 | 0.66 | 0.12 | Body: 6; Animal: 8; Blood: 4; Object: 1; People: 1; Fire: 4 |
| **NFT run3** | 6.02 | 0.57 | 3.01 | 0.88 | 0.63 | 0.11 | Body: 8; Animal: 8; Blood: 3; Object: 2; Fire: 3 |
| **NFT run4** | 6.00 | 0.52 | 3.13 | 0.73 | 0.64 | 0.12 | Body: 6; Animal: 8; Blood: 4; Object: 2; Fire: 4 |
| **Transfer** | 6.01 | 0.65 | 3.13 | 0.75 | 0.64 | 0.10 | Body: 5; Animal: 9; Blood: 4; Object: 2; Fire: 4 |

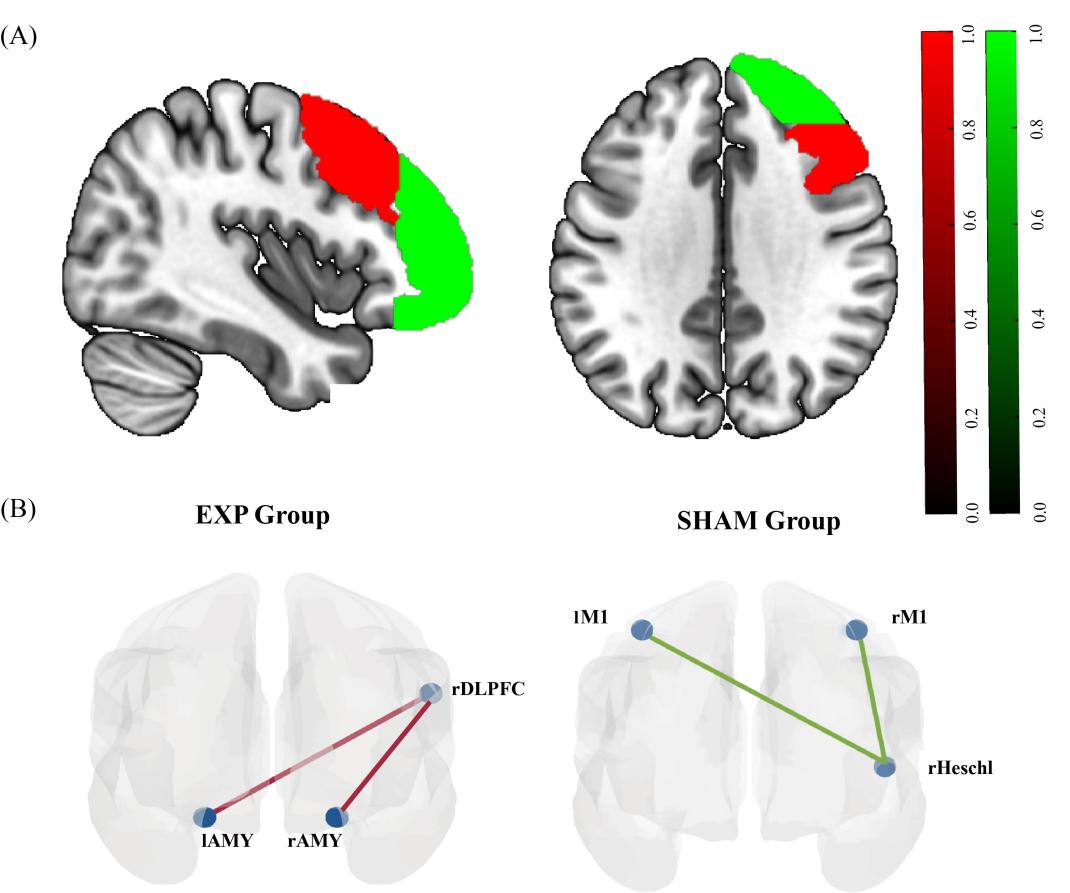

**Fig. S1 Mask and training circuits**

(A) The mask incorporate right frontal pole (shown in green) and right middle frontal gyrus (shown in red) from the Harvard Oxford atlas. This mask was used during the localizer run to help identify the peak-activated coordinate of the right DLPFC ROI, which was subsequently used for the active NFT. (B) Online training circuit for EXP group and SHAM group during NFT.

###### Task-evoked activation results

For report completeness, we also examined task-evoked activation at the whole brain level in the three tasks. In the LocT, group-level analyses revealed an extensive activation pattern similarly between the EXP and SHAM groups in (i) the cognitive control and emotional regulation networks including the bilateral DLPFC extending to the bilateral inferior prefrontal gyrus (IFG), bilateral thalamus (Thal) and bilatAMY, (ii) the learning and memory networks including the bilateral hippocampus, bilateral putamen (PUT) extending to the bilateral globus pallidus (GP), and (iii) the visual information processing and integration regions including the bilateral inferior occipital gyrus (IOG) extending to the bilateral middle occipital gyrus (MOG) and the right MTG (*p* < 0.01, FDR-corrected; Fig. S2A). In the NFT, a similar activation pattern to the LocT was found for both the two groups, except that activation in the rAMY was only found in the SHAM group and more regions in the learning system (e.g., caudate and GP) were found for both groups (*p* < 0.01, FDR-corrected; Fig. S2B). In the TransT run, the SHAM group exhibited a similar activation pattern to those observed in the LocT, while the EXP group did not show significant activation in the bilatAMY and INS involved in negative emotional processing (*p* < 0.01, FDR-corrected; Fig. S2C), which were observed in the SHAM group and in the LocT. No significant group differences were found for all the three tasks with the same threshold (*p* < 0.01, FDR-corrected).

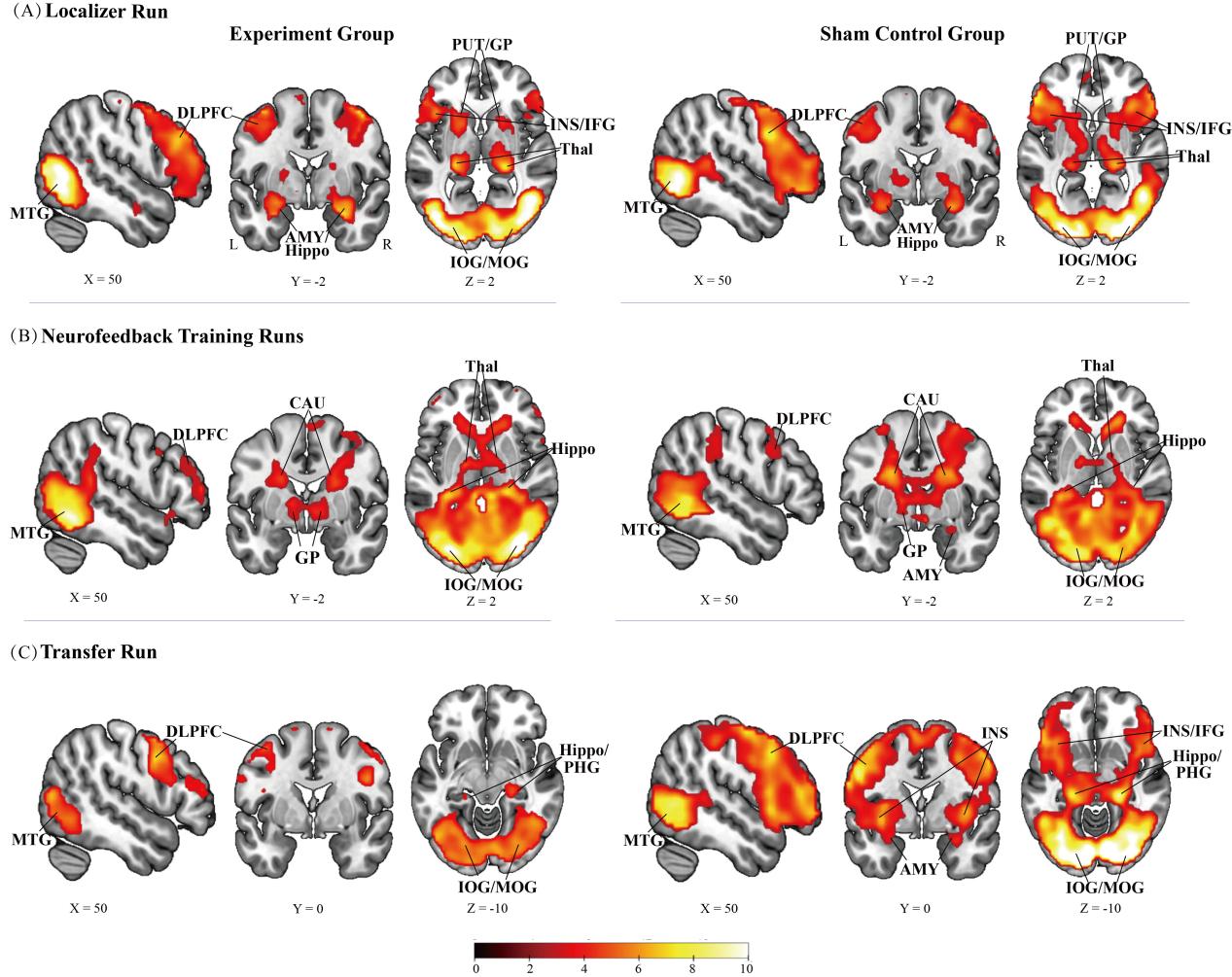

**Fig. S2 Whole-brain activation results**

(A) Whole-brain activation patterns in the LocT for the EXP and SHAM groups respectively. (B) Whole-brain activation patterns in the NFT for the EXP and SHAM groups respectively. (C) Whole-brain activation patterns in the TransT for the EXP and SHAM groups respectively. All results are presented with a threshold of *p* < 0.01, FDR-corrected. SFG: superior frontal gyrus; MTG: Middle Temporal gyrus; CAU: caudate; Thal: thalamus; Hippo: hippocampus gyrus; PHG: parahippocampal gyrus; AMY: amygdala; INS: insula; IFG: inferior frontal gyrus; PUT: putamen; GP: globus pallidus; IOG: inferior occipital gyrus; MOG: middle occipital gyrus; Error bars represent standard error, white rectangular represents the mean value.

**Table S4.**

Whole-brain activation results in the LocT for the EXP and SHAM groups respectively.

| Brain region | BA | No.Voxels | Peak t-value | X | Y | Z |
| --- | --- | --- | --- | --- | --- | --- |
| **EXP_LocT** |  |  |  |  |  |  |
| L. Fusiform | 19 | 11587 | 15.44 | -24 | -70 | -13 |
| Lingual Gyrus | 18 |  | 14.71 | 9 | -79 | -10 |
| Cerebellum Posterior Lobe |  |  | 13.5 | -6 | -82 | -13 |
| Middle Occipital Gyrus |  |  | 12.21 | -36 | -85 | -1 |
| Fusiform |  |  | 11.77 | 27 | -73 | -13 |
| Inferior Occipital Gyrus | 19 |  | 11.65 | -42 | -73 | -10 |
| Inferior Occipital Gyrus |  |  | 11.23 | 39 | -76 | -1 |
| Middle Occipital Gyrus | 19 |  | 10.99 | 48 | -73 | 5 |
| Dorsolateral Prefrontal Cortex | 46 |  | 8.83 | 54 | 29 | 29 |
| Parahippocampus |  |  | 8.8 | -18 | -34 | -7 |
| Amygdala |  |  | 8.17 | 27 | -1 | -22 |
| Amygdala |  |  | 7.6 | -27 | -4 | -22 |
| Thalamus |  |  | 7.34 | 21 | -28 | 2 |
| Hippocampus |  |  | 7.25 | 21 | -7 | -19 |
| Hippocampus |  |  | 7.02 | -27 | -7 | -13 |
| Insula |  |  | 6.64 | -27 | 17 | -4 |
| Inferior Frontal Gyrus |  |  | 6.38 | -45 | 20 | 20 |
| Dorsolateral Prefrontal Cortex | 8/9 |  | 6.28 | -48 | 20 | 41 |
| Putamen |  |  | 5.29 | -18 | 11 | 2 |
| Putamen |  |  | 4.93 | 24 | 11 | -1 |
| Thalamus |  |  | 4.29 | -21 | -28 | 5 |
| R.Superior Temporal Gyrus |  | 28 | 4.74 | 60 | -40 | 14 |
| L.Ventromedial Prefrontal Cortex | 11 | 17 | 4.25 | -3 | 56 | -13 |
| **SHAM_LocT** |  |  |  |  |  |  |
| L. Calcarine |  | 12578 | 16.68 | -6 | -91 | -7 |
| Middle Occipital Gyrus |  |  | 14.48 | 27 | -94 | 5 |
| Middle Occipital Gyrus |  |  | 13.85 | 33 | -88 | 5 |
| Hippocampus | 27 |  | 11.52 | 21 | -31 | -4 |
| Middle Occipital Gyrus |  |  | 11.26 | -30 | -91 | 5 |
| Middle Temporal Gyrus | 37 |  | 10.98 | 48 | -61 | -1 |
| Parahippocampus |  |  | 8.94 | -18 | -31 | -7 |
| Hippocampus | 27 |  | 8.15 | -18 | -31 | -4 |
| Dorsolateral Prefrontal Cortex | 8 |  | 7.43 | -54 | 8 | 41 |
| Inferior Frontal Gyrus | 47 |  | 6.8 | -45 | 23 | -13 |
| Dorsolateral Prefrontal Cortex | 9 |  | 6.48 | 57 | 14 | 38 |
| Inferior Frontal Gyrus |  |  | 6.35 | 51 | 20 | 23 |
| Amygdala |  |  | 5.17 | 27 | -4 | -16 |
| Amygdala |  |  | 5.11 | -27 | -4 | -16 |
| Putamen |  |  | 4.83 | -21 | 5 | 8 |
| Insula | 47 |  | 4.7 | -27 | 14 | -19 |
| Thalamus |  |  | 4.6 | 12 | -16 | 2 |
| Insula |  |  | 4.52 | 27 | 17 | -19 |
| Putamen |  |  | 4.4 | 21 | 5 | 5 |
| Thalamus |  |  | 4.35 | -12 | -10 | -1 |
| L. SupraMarginal Gyrus |  | 23 | 3.72 | -51 | -43 | 23 |
| SupraMarginal Gyrus | 40 |  | 3.02 | -60 | -46 | 32 |

All results were reported with a threshold of *p* < 0.01 FDR-corrected.

**Table S5.**

Whole-brain activation results in the NFT for the EXP and SHAM groups respectively.

| Brain region | BA | No.Voxels | Peak t-value | X | Y | Z |
| --- | --- | --- | --- | --- | --- | --- |
| **EXP_NFT** |  |  |  |  |  |  |
| L. Superior Parietal Lobule | 7 | 17600 | 11.21 | 27 | -64 | 50 |
| Middle Occipital Gyrus |  |  | 10.74 | -27 | -88 | 3 |
| Middle Occipital Gyrus |  |  | 10.58 | 30 | -79 | 17 |
| Inferior Temporal Gyrus | 37 |  | 9.18 | 51 | -55 | -10 |
| Middle Temporal Gyrus |  |  | 8.11 | 48 | -58 | 2 |
| Dorsolateral Prefrontal Cortex |  |  | 5.65 | -42 | 47 | 23 |
| Dorsolateral Prefrontal Cortex |  |  | 5.59 | 36 | 53 | 26 |
| Thalamus |  |  | 5.38 | -15 | -7 | 5 |
| Dorsolateral Prefrontal Cortex | 46 |  | 5.31 | 54 | 41 | 8 |
| Dorsolateral Prefrontal Cortex | 10 |  | 5.17 | -42 | 50 | 14 |
| Caudate |  |  | 4.98 | 18 | 14 | 20 |
| Dorsolateral Prefrontal Cortex | 10 |  | 4.73 | 39 | 56 | 17 |
| Dorsolateral Prefrontal Cortex | 10 |  | 4.53 | -45 | 50 | 5 |
| Caudate |  |  | 4.44 | -18 | 2 | 23 |
| Globus Pallidus |  |  | 4.36 | 9 | -1 | -1 |
| Superior Temporal Gyrus |  |  | 4.09 | 54 | 17 | -10 |
| Globus Pallidus |  |  | 4.09 | -15 | -4 | 2 |
| Superior Temporal Gyrus | 38 |  | 4.02 | 57 | 11 | -16 |
| Thalamus |  |  | 3.68 | 15 | -10 | 11 |
| Inferior Frontal Gyrus |  |  | 3.36 | 57 | 17 | -1 |
| **SHAM_NFT** |  |  |  |  |  |  |
| R. Inferior Temporal Gyrus | 18526 |  | 9.94 | 42 | -49 | -4 |
| Middle Occipital Gyrus |  |  | 9.72 | -30 | -73 | -1 |
| Middle Occipital Gyrus |  |  | 9.34 | 33 | -67 | 35 |
| Hypothalamus |  |  | 5.81 | 6 | -1 | -10 |
| Thalamus |  |  | 5.58 | 21 | -28 | 8 |
| Caudate |  |  | 5.56 | -18 | -4 | 23 |
| Caudate |  |  | 5.28 | 18 | -4 | 23 |
| Hippocampus |  |  | 5.24 | 39 | -28 | -10 |
| Globus Pallidus |  |  | 5.2 | -18 | -10 | -7 |
| Amygdala |  |  | 5 | 27 | 2 | -19 |
| Dorsolateral Prefrontal Cortex |  |  | 4.72 | 39 | 44 | 32 |
| Dorsolateral Prefrontal Cortex | 46 |  | 4.01 | -54 | 29 | 23 |
| Globus Pallidus |  |  | 3.94 | 24 | -13 | -7 |
| Hypothalamus |  |  | 3.78 | -3 | -1 | -7 |
| Precental Gyrus |  |  | 3.55 | -36 | -22 | 65 |

All results were reported with a threshold of *p* < 0.01 FDR-corrected.

**Table S6.**

Whole-brain activation results in the TransT for the EXP and SHAM groups respectively.

| Brain region | BA | No.Voxels | Peak t-value | X | Y | Z |
| --- | --- | --- | --- | --- | --- | --- |
| **EXP_TransT** |  |  |  |  |  |  |
| R. Fusiform Gyrus |  | 3626 | 8.3 | 33 | -49 | -22 |
| Cerebellum Posterior Lobe |  |  | 7.85 | 24 | -73 | -19 |
| Cerebellum Posterior Lobe | 18 |  | 7.44 | -15 | -85 | -19 |
| R. Inferior Frontal Gyrus |  | 470 | 7.96 | 48 | 5 | 35 |
| Inferior Frontal Gyrus |  |  | 6.97 | 54 | 14 | 38 |
| Dorsolateral Prefrontal Cortex | 8 |  | 6.88 | 54 | 8 | 44 |
| Dorsolateral Prefrontal Cortex |  |  | 6.23 | 48 | 2 | 53 |
| L. Superior Parietal Lobule | 7 | 121 | 5.7 | -21 | -64 | 62 |
| Superior Parietal Lobule | 7 |  | 4.86 | -18 | -67 | 47 |
| Superior Parietal Lobule | 7 |  | 4.7 | -24 | -55 | 47 |
| Middle Temporal Gyrus | 37 |  | 7.02 | 45 | -70 | 5 |
| L. Dorsolateral Prefrontal Cortex | 6 | 70 | 5.65 | -39 | -1 | 59 |
| Dorsolateral Prefrontal Cortex |  |  | 4.05 | -36 | -1 | 41 |
| Dorsolateral Prefrontal Cortex |  |  | 4.02 | -54 | 2 | 44 |
| L. Inferior Frontal Gyrus | 9 | 140 | 5.4 | -57 | 8 | 35 |
| Inferior Frontal Gyrus |  |  | 4.65 | -39 | 5 | 20 |
| Inferior Frontal Gyrus |  |  | 4.36 | -45 | 23 | 17 |
| Dorsolateral Prefrontal Cortex |  |  | 4.08 | -48 | 29 | 17 |
| R. Supplementary Motor Area | 6 | 25 | 4.79 | 12 | 5 | 68 |
| Supplementary Motor Area |  |  | 4.03 | 15 | 14 | 65 |
| L. Hippocampus | 27 | 15 | 4.3 | -21 | -31 | -4 |
| L. Superior Frontal Gyrus | 6 | 16 | 4.17 | -18 | -1 | 71 |
| Supplementary Motor Area |  |  | 4.09 | -12 | 11 | 68 |
| L. Superior Frontal Gyrus | 6 | 14 | 4.01 | -6 | 8 | 56 |
| **SHAM_TransT** |  |  |  |  |  |  |
| L. Inferior Occipital Gyrus | 18 | 18330 | 19.44 | -30 | -88 | -4 |
| Middle Occipital Gyrus |  |  | 16.9 | 30 | -94 | 5 |
| Lingual Gyrus |  |  | 15.85 | -9 | -88 | -10 |
| Hippocampus | 27 |  | 12.43 | 21 | -31 | -4 |
| Hippocampus | 27 |  | 9.73 | -18 | -34 | -4 |
| Parahippocampus |  |  | 9.16 | 21 | -34 | -7 |
| Middle Temporal Gyrus |  |  | 8.39 | 51 | -58 | 5 |
| Inferior Frontal Gyrus |  |  | 7.77 | -54 | 14 | 23 |
| Insula |  |  | 7.03 | -36 | 2 | -4 |
| Parahippocampus |  |  | 6.94 | -18 | -37 | -7 |
| Insula |  |  | 6.47 | 39 | -1 | 2 |
| Amygdala |  |  | 4.59 | -24 | -4 | -19 |
| Amygdala |  |  | 4.42 | 30 | -4 | -19 |
| L. Superior Frontal Gyrus |  | 24 | 4.15 | -18 | 47 | 41 |
| Superior Frontal Gyrus |  |  | 3.23 | -9 | 53 | 38 |

All results were reported with a threshold of *p* < 0.01 FDR-corrected.

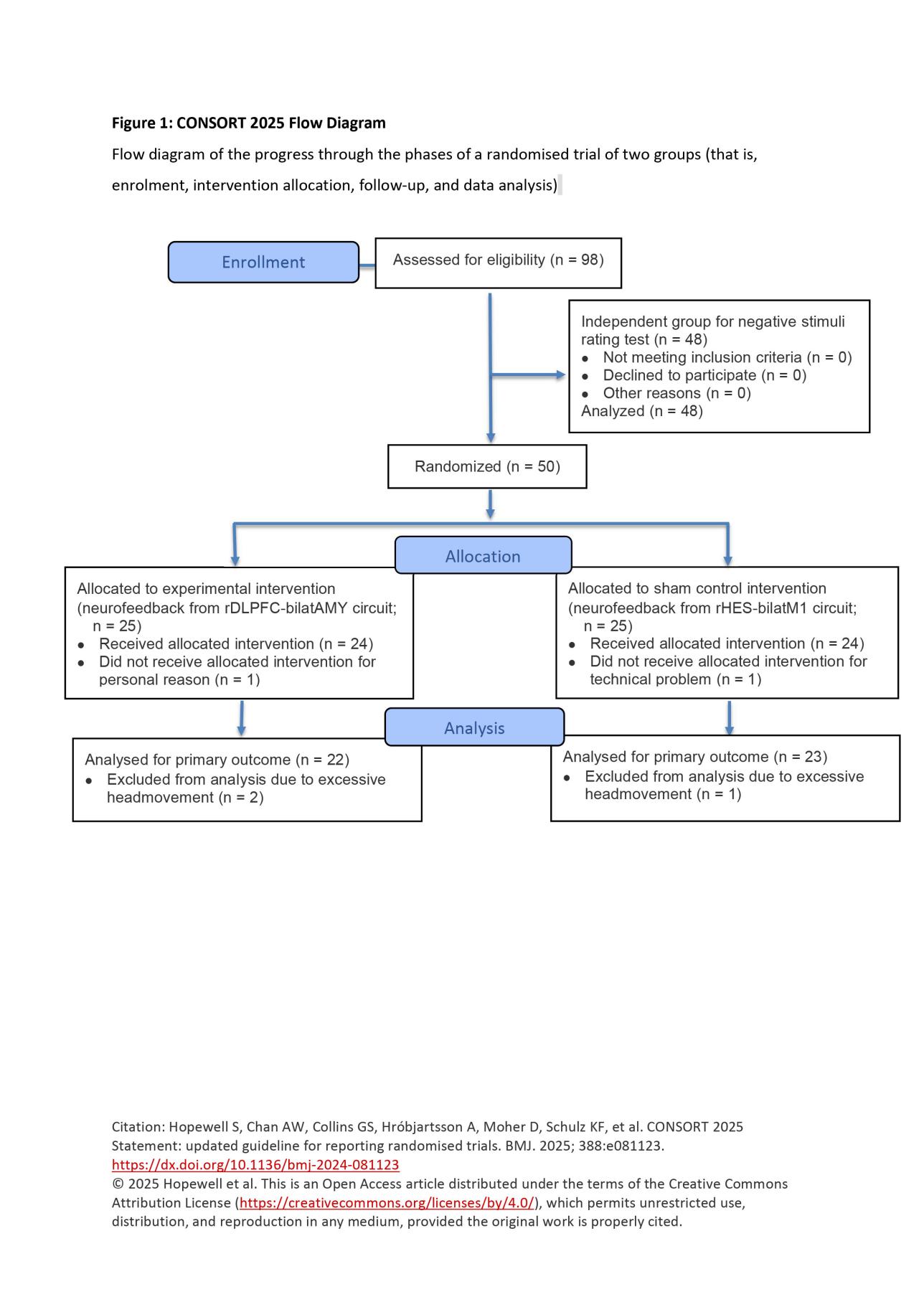

**Fig.S3 CONSORT 2025 flow diagram for experimemt**

**Table S7.**

CONSORT checklist for functional connectivity-based neurofeedback of the DLPFC-amygdala pathway.

|  | Section/topic | No | CONSORT 2025 checklist item description | Reported on page no. |
| --- | --- | --- | --- | --- |
|  | **Title and abstract** | | |  |
|  | Title and structured abstract | 1a | Identification as a randomised trial | 2 |
|  |  | 1b | Structured summary of the trial design, methods, results, and conclusions | 2 |
|  | **Open science** | | |  |
|  | Trial registration | 2 | Name of trial registry, identifying number (with URL) and date of registration | 5 |
|  | Protocol and statistical analysis plan | 3 | Where the trial protocol and statistical analysis plan can be accessed | 4-8 |
|  | Data sharing | 4 | Where and how the individual de-identified participant data (including data dictionary), statistical code and any other materials can be accessed | N/A |
|  | Funding and conflicts of interest | 5a | Sources of funding and other support (eg, supply of drugs), and role of funders in the design, conduct, analysis and reporting of the trial | 14 |
|  |  | 5b | Financial and other conflicts of interest of the manuscript authors | 14 |
|  | **Introduction** | | |  |
|  | Background and rationale | 6 | Scientific background and rationale | 2-4 |
|  | Objectives | 7 | Specific objectives related to benefits and harms | 3-4 |
|  | **Methods** | | |  |
|  | Patient and public involvement | 8 | Details of patient or public involvement in the design, conduct and reporting of the trial | 4-5 |
|  | Trial design | 9 | Description of trial design including type of trial (eg, parallel group, crossover), allocation ratio, and framework (eg, superiority, equivalence, non-inferiority, exploratory) | 4-5 |
|  | Changes to trial protocol | 10 | Important changes to the trial after it commenced including any outcomes or analyses that were not prespecified, with reason | N/A |
|  | Trial setting | 11 | Settings (eg, community, hospital) and locations (eg, countries, sites) where the trial was conducted | 4 |
|  | Eligibility criteria | 12a | Eligibility criteria for participants | 4 |
|  |  | 12b | If applicable, eligibility criteria for sites and for individuals delivering the interventions (eg, surgeons, physiotherapists) | N/A |
|  | Intervention and comparator | 13 | Intervention and comparator with sufficient details to allow replication. If relevant, where additional materials describing the intervention and comparator (eg, intervention manual) can be accessed | 4-6 |
|  | Outcomes | 14 | Prespecified primary and secondary outcomes, including the specific measurement variable (eg, systolic blood pressure), analysis metric (eg, change from baseline, final value, time to event), method of aggregation (eg, median, proportion), and time point for each outcome | 5-8 |
|  | Harms | 15 | How harms were defined and assessed (eg, systematically, non-systematically) | N/A |
|  | Sample size | 16a | How sample size was determined, including all assumptions supporting the sample size calculation | 4 |
|  |  | 16b | Explanation of any interim analyses and stopping guidelines | N/A |
|  | Randomisation: |  |  |  |
|  | Sequence generation | 17a | Who generated the random allocation sequence and the method used | N/A |
|  |  | 17b | Type of randomisation and details of any restriction (eg, stratification, blocking and block size) | N/A |
|  |  |  |  | **Reported on page no.** |
|  | Allocation concealment mechanism | 18 | Mechanism used to implement the random allocation sequence (eg, central computer/telephone; sequentially numbered, opaque, sealed containers), describing any steps to conceal the sequence until interventions were assigned | N/A |
|  | Implementation | 19 | Whether the personnel who enrolled and those who assigned participants to the interventions had access to the random allocation sequence | 4 |
|  | Blinding | 20a | Who was blinded after assignment to interventions (eg, participants, care providers, outcome assessors, data analysts) | N/A |
|  |  | 20b | If blinded, how blinding was achieved and description of the similarity of interventions | N/A |
|  | Statistical methods | 21a | Statistical methods used to compare groups for primary and secondary outcomes, including harms | 5 |
|  |  | 21b | Definition of who is included in each analysis (eg, all randomised participants), and in which group | 4-5 |
|  |  | 21c | How missing data were handled in the analysis | N/A |
|  |  | 21d | Methods for any additional analyses (eg, subgroup and sensitivity analyses), distinguishing prespecified from post hoc | N/A |
|  | **Results** | | |  |
|  | Participant flow, including flow diagram | 22a | For each group, the numbers of participants who were randomly assigned, received intended intervention, and were analysed for the primary outcome | 8-13 |
|  |  | 22b | For each group, losses and exclusions after randomisation, together with reasons | 4-5 |
|  | Recruitment | 23a | Dates defining the periods of recruitment and follow-up for outcomes of benefits and harms | 5 |
|  |  | 23b | If relevant, why the trial ended or was stopped | N/A |
|  | Intervention and comparator delivery | 24a | Intervention and comparator as they were actually administered (eg, where appropriate, who delivered the intervention/comparator, how participants adhered, whether they were delivered as intended (fidelity)) | N/A |
|  |  | 24b | Concomitant care received during the trial for each group | N/A |
|  | Baseline data | 25 | A table showing baseline demographic and clinical characteristics for each group | 9 |
|  | Numbers analysed,  outcomes and estimation | 26 | For each primary and secondary outcome, by group:  ● the number of participants included in the analysis  ● the number of participants with available data at the outcome time point  ● result for each group, and the estimated effect size and its precision (such as 95% confidence interval)  ● for binary outcomes, presentation of both absolute and relative effect size | 8-13 |
|  | Harms | 27 | All harms or unintended events in each group | N/A |
|  | Ancillary analyses | 28 | Any other analyses performed, including subgroup and sensitivity analyses, distinguishing pre-specified from post hoc | N/A |
|  | **Discussion** | | | 13-14 |
|  | Interpretation | 29 | Interpretation consistent with results, balancing benefits and harms, and considering other relevant evidence | 14-15 |
|  | Limitations | 30 | Trial limitations, addressing sources of potential bias, imprecision, generalisability, and, if relevant, multiplicity of analyses |  |

**Table S8.**

Consensus on the Reporting and Experimental Design of clinical and cognitive-behavioural Neurofeedback studies (CRED-nf) best practices checklist 2020*

| **Domain** | **Item #** | **Checklist item** | **Reported on page #** |
| --- | --- | --- | --- |
| **Pre-experiment** | | | |
|  | 1a | Pre-register experimental protocol and planned analyses | 5 |
|  | 1b | Justify sample size | 4-5 |
| **Control groups** | | | |
|  | 2a | Employ control group(s) or control condition(s) | 6 |
|  | 2b | When leveraging experimental designs where a double-blind is possible, use a double-blind | 4-5 |
|  | 2c | Blind those who rate the outcomes, and when possible, the statisticians involved | N/A |
|  | 2d | Examine to what extent participants and experimenters remain blinded | N/A |
|  | 2e | In clinical efficacy studies, employ a standard-of-care intervention group as a benchmark for improvement | N/A |
| **Control measures** | | | |
|  | 3a | Collect data on psychosocial factors | 5 |
|  | 3b | Report whether participants were provided with a strategy | 5 |
|  | 3c | Report the strategies participants used | 5 |
|  | 3d | Report methods used for online-data processing and artifact correction | 7 |
|  | 3e | Report condition and group effects for artifacts | 9-10 |
| **Feedback specifications** | | | |
|  | 4a | Report how the online-feature extraction was defined | 6-7 |
|  | 4b | Report and justify the reinforcement schedule | 6 |
|  | 4c | Report the feedback modality and content | 5 |
|  | 4d | Collect and report all brain activity variable(s) and/or contrasts used for feedback, as displayed to experimental participants | 7-8 |
|  | 4e | Report the hardware and software used | 7-8 |
| **Outcome measures** | | | |
| Brain | 5a | Report neurofeedback regulation success based on the feedback signal | 9-10 |
|  | 5b | Plot within-session and between-session regulation blocks of feedback variable(s), as well as pre-to-post resting baselines or contrasts | 10 |
|  | 5c | Statistically compare the experimental condition/group to the control condition(s)/group(s) (not only each group to baseline measures) | 9-10 |
| Behaviour | 6a | Include measures of clinical or behavioural significance, defined a priori, and describe whether they were reached | 8-9 |
|  | 6b | Run correlational analyses between regulation success and behavioural outcomes | N/A |
| **Data storage** | | |  |
|  | 7a | Upload all materials, analysis scripts, code, and raw data used for analyses, as well as final values, to an open access data repository, when feasible | N/A |
